# Metabolic enzyme cost explains variable trade-offs between microbial growth rate and yield

**DOI:** 10.1101/111161

**Authors:** Meike T. Wortel, Elad Noor, Michael Ferris, Frank J. Bruggeman, Wolfram Liebermeister

**Author notes:** These authors contributed equally to this work (Meike T. Wortel, Elad Noor).

## Abstract

Microbes may maximize the number of daughter cells per time or per amount of nutrients consumed. These two strategies correspond, respectively, to the use of enzyme-efficient or substrate-efficient metabolic pathways. In reality, fast growth is often associated with wasteful, yield-inefficient metabolism, and a general thermodynamic trade-off between growth rate and biomass yield has been proposed to explain this. We studied growth rate/yield trade-offs by using a novel modeling framework, Enzyme-Flux Cost Minimization (EFCM) and by assuming that the growth rate depends directly on the enzyme investment per rate of biomass production. In a comprehensive mathematical model of core metabolism in *E. coli*, we screened all elementary flux modes leading to cell synthesis, characterized them by the growth rates and yields they provide, and studied the shape of the resulting rate/yield Pareto front. By varying the model parameters, we found that the rate/yield trade-off is not universal, but depends on metabolic kinetics and environmental conditions. A prominent trade-off emerges under oxygen-limited growth, where yield-inefficient pathways support a 2-to-3 times higher growth rate than yield-efficient pathways. EFCM can be widely used to predict optimal metabolic states and growth rates under varying nutrient levels, perturbations of enzyme parameters, and single or multiple gene knockouts.

**Author Summary:** When cells compete for nutrients, those that grow faster and produce more offspring per time are favored by natural selection. In contrast, when cells need to maximize the cell number at a limited nutrient supply, fast growth does not matter and an efficient use of nutrients (i.e. high biomass yield) is essential. This raises a basic question about metabolism: can cells achieve high growth rates and yields simultaneously, or is there a conflict between the two goals? Using a new modeling method called Enzymatic Flux Cost Minimization (EFCM), we predict cellular growth rates and find that growth rate/yield trade-offs and the ensuing preference for enzyme-efficient or substrate-efficient metabolic pathways are not universal, but depend on growth conditions such as external glucose and oxygen concentrations.

## Introduction

Metabolic networks are shaped by evolution. In well-mixed, nutrient-rich environments, fast-growing bacteria are favored by natural selection. Such environments are commonly studied in laboratory settings, but natural environments are more diverse. In isolated ecological niches with limited resources, it is the total number of offspring cells, rather than fast growth, that determines evolutionary success. This puts a selection pressure on biomass yield (biomass produced per amount of the limiting nutrient, e.g. glucose) rather than on growth rate (biomass produced per time and per cell biomass). Mechanistically, growth rate and yield might be expected to go hand in hand. It seems logical that a cell with a higher yield – i.e. one that can produce offspring from a smaller amount of nutrients – would also produce a larger number of offspring per time. However, in experiments we observe exactly the opposite; many fast-growing cells employ low-yield metabolic pathways (e.g. yeast cells (Crabtree effect) and cancer cells (Warburg effect)[1]), and also many bacteria display a wasteful respiro-fermentative overflow metabolism and still attain high growth rates. Pure respiratory growth would give rise to a higher biomass yield per mole of glucose, but to lower growth rates.

Since yield-inefficient metabolic strategies are widely observed, under various circumstances and in evolutionarily unrelated organisms, it has been suggested that growth rate and yield may be in conflict for physicochemical reasons. During evolution, such a conflict may lead to “tragedy-of-the-commonsd” situations in which yield-inefficient microbes gain an evolutionary advantage by over-exploiting shared resources [2–4]. The hypothesis of a general trade-off is supported by simple cell models in which high-yield pathways display lower thermodynamic forces or higher enzyme costs [5–7].

The rate/yield trade-off has been tested by lab-evolution experiments with fast-growing microorganisms, with varying levels of success. Growth rate and yield have been compared between different wild-type and evolved microbial strains [8–11], but most studies found poor correlations between growth rate and yield. Novak et al. [9] found a negative correlation within evolved *E. coli* populations, indicating a rate/yield trade-off. A rare example of bacteria evolving for high yield in the laboratory was in the work of Bachmann et al. [12]. In their protocol, cells grow in separate droplets in a medium-in-oil suspension, simulating a fragmented environment, and offspring cells are mixed when the nutrients in the droplets have been depleted, and then resuspended. This creates a strong selection pressure for maximizing biomass yield. Indeed, the strains evolved towards higher yields at the expense of their growth rate, again indicating a trade-off between the two objectives. However, evidence from all these experiments may not be conclusive, because microorganisms may behave sub-optimally in the laboratory experiments.

Thus, is the rate/yield trade-off universal? We claim that the answer to this question lies in metabolism, especially in enzyme demand. At balanced growth, the relative amounts of all cell components remain constant in time, including the protein fraction associated with metabolic enzymes. If a metabolic strategy achieves a given biomass synthesis rate at a lower enzyme demand, the freed protein resources can be reallocated to other cellular processes that contribute to growth, and the cell’s growth rate can increase. Thus, a metabolic strategy will be growth-optimal if it minimizes enzyme cost at a given biomass synthesis rate [13].

In theory, the use of a high-yield flux mode affects the growth rate in two opposite ways. On the one hand, a high-yield mode achieves the same rate of biomass production at a lower glycolytic rate, and the lower enzyme demand in glycolysis allows for a higher growth rate. On the other hand, high-yield modes dissipate less Gibbs free energy [5], which may slow down the reactions and must be compensated by higher enzyme levels, leading to lower growth rates [7, 14, 15]. The second effect may be obscured if another substrate, such as oxygen, provides additional driving force.

When the first effect dominates, high-yield modes allow for a higher biomass production per enzyme invested, so yield and growth rate are maximized by a single flux mode. When the second effect dominates, it is low-yield modes that provide a growth advantage [6, 13, 16–18], and there will be a trade-off: growth rate and yield are maximized by different flux modes, and there may be other modes in between that provide optimal compromises. In summary, a rate/yield trade-off in cells reflects a trade-off between *enzyme efficiency* and *substrate efficiency* in metabolism; and since the enzyme cost of a given pathway flux depends on external conditions, the occurrence of rate/yield trade-off will be condition-dependent as well.

How can we describe this by models? The specific growth rate *μ* for exponentially growing cells is given by the rate of biomass synthesis per cell dry weight and is typically measured in grams of biomass per gram cell dry weight per hour. The biomass yield*Y*_*X/S*_ is measured in grams of biomass per carbon mole of nutrient (i.e. per 1/6 mole of glucose). If the carbon uptake rate *v*_*S*_ were known, we could directly convert between yield and growth rate using this formula: 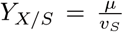. However, since carbon uptake, yield, and growth rate are tightly coupled, the changes in *v*_*S*_ are hard to predict. Classical Flux Balance Analysis (FBA) places an upper bound on *v*_*S*_. If this is the only active flux bound, then maximizing biomass rate coincides with maximizing biomass yield, leaving no possibility for rate/yield trade-offs. Other constraint-based methods, such FBA with Molecular Crowding [19] or Resource Balance Analysis [20], account for enzyme costs. They can be used to explore the trade-off, but they are not fully quantitative because they ignore the kinetic and thermodynamic effects of varying metabolite concentrations (see Discussion section for details).

In [21], a kinetic pathway model was used to directly compute the enzyme costs. Two variants of glycolysis, both common among bacteria, were compared by their ATP yields on glucose and by their ATP production per enzyme investment. At a given glucose influx, the Embden-Meyerhof-Parnas (EMP) pathway yields twice as much ATP, but was found to use more than 4 times as much enzyme than the Entner-Doudoroff (ED) pathway. This suggested that cells under yield selection should use the EMP pathway, while cells under rate selection should use the ED pathway instead. Aside from simple approximations [22, 23], the enzyme economics of other metabolic choices, e.g. respiration *versus* fermentation, and the resulting trade-offs, remain to be quantified.

Here we combine a calculation of enzyme cost, based on kinetic models, with elementary flux mode analysis. Elementary flux modes (EFMs) describe the fundamental ways in which a metabolic network can operate [24–27]. Among the steady-state flux modes, EFMs are minimal in the sense that they do not contain any smaller subnetworks that can support a steady-state flux mode [24, 25, 27]. EFMs might be expected to have simple shapes in the network, but since biomass production requires many different precursors, biomass-producing EFMs can be highly branched. All biomass-producing EFMs are free of thermodynamically infeasible loops, and if the flux directions are predefined, the set of steady-state flux distribution is a convex polytope spanned by the EFMs. The EFMs of a metabolic network can be enumerated, and thermodynamically infeasible modes can be efficiently discarded [28, 29], but in practice an enumeration of EFMs may be impossible because of their large number. EFMs have a remarkable property, which makes them well-suited for studying rate/yield trade-offs: in kinetic metabolic models, the biomass production per enzyme investment is maximized by a vertex point of the flux polytope, and in models without flux bounds, all these vertices are EFMs [30–32]. The yield of an EFM, defined as the output flux divided by the input flux, is easy to compute and it is again an EFM that achieves the maximal yield among flux modes. Therefore, to find flux modes that maximize cell growth, we can enumerate the EFMs and assess them one by one; and to determine rate/yield trade-offs, we simply plot yields versus growth rates of all EFMs (Figure 1(a)).

**Figure 1:**
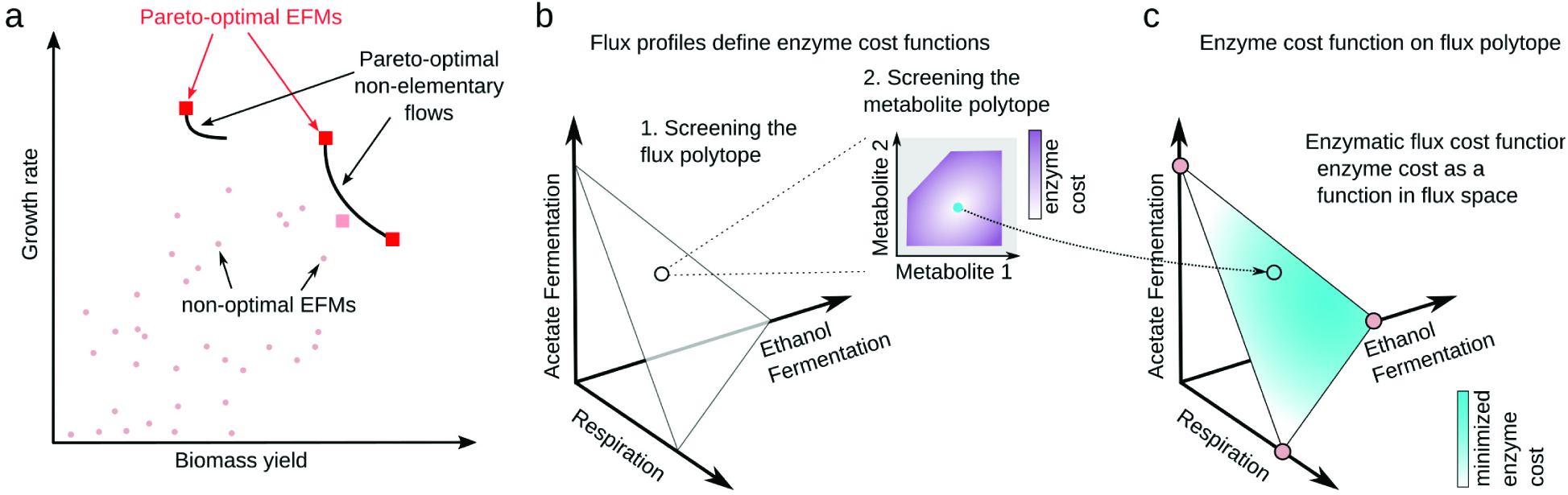
Rate/yield trade-offs and calculation of growth-optimal fluxes. (a) Rate/yield spectrum of Elementary Flux Modes (EFMs) (schematic drawing). In the scatter plot, EFMs are represented by points indicating biomass yield and maximal achievable growth rate in a given simulation scenario. Pareto-optimal EFMs are marked by red squares. The set of Pareto-optimal flux modes (black lines) contains also non-elementary flux modes. An EFM may be Pareto-optimal when compared to other EFMs, but not when compared to all possible flux modes (e.g. the EFM below the Pareto front marked by a the pink square). Growth rate and yield are positively correlated in the entire point cloud, but the points along the Pareto front show a negative correlation, indicating a trade-off. (b) Enzyme cost of metabolic fluxes. The space of stationary flux distributions is spanned by three EFMs (hypothetical example). The flux modes, scaled to unit biomass production, form a triangle. To compute the enzyme cost of a flux mode, we determine the optimal enzyme and metabolite levels. To do so, we minimize the enzymatic cost on the metabolite polytope (inset graphics) by solving a convex optimality problem called Enzyme Cost Minimization (ECM). (c) Calculation of optimal flux modes. The enzymatic cost is a concave function on the flux polytope, and its optimal points must be polytope vertices. In models without flux bounds, these vertices are EFMs and optimal flux modes can be found by screening all EFMs and choosing the one with the minimal cost.

## Results

### Computing the cell growth rate

To predict optimal metabolic fluxes and cell growth rates, we developed Enzyme-Flux Cost Minimization (EFCM), a method for computing flux modes that realize a linear flux objective at a minimal enzyme cost. Constraint-based methods such as Flux Balance Analysis are entirely based on reaction stoichiometries. Some of them also use approximate enzyme costs, for instance the sum of absolute fluxes [33] or other linear/quadratic functions of the flux vector [19]. EFCM, in contrast, computes enzyme cost based on a given kinetic model. In our model, the flux objective represents biomass production, i.e. the production of small molecules and macromolecules that constitute the cell and do not explicitly appear in the network model. Below we argue that enzyme-optimal flux modes, with such a flux objective, are the ones that allow for maximal growth rates.

To compute the maximal growth rate achievable, we use a kinetic model of metabolism, consider all possible flux modes, and compute for of them the optimal enzyme allocation pattern, i.e. the pattern that realizes the required fluxes at a minimal total enzyme investment. Enzyme Cost Minimization (ECM) is a method that finds optimal enzyme and metabolite profiles supporting a given flux distribution [34]. The ECM problem can be quickly solved using convex optimization, and the minimal enzyme cost of all EFMs can be computed in reasonable time (a few minutes on a shared server, for models with ~ 10^3^ EFMs such as *E. coli* core metabolism). Knowing the enzyme investment per biomass production, we next compute the cellular growth rate. For each EFM, the enzyme demand per biomass production is translated into a mass doubling time (i.e. the amount of time that metabolism would have to run in order to duplicate all metabolic enzymes assumed in our model). The mass doubling time can be translated into a cell growth rate by a semi-empirical formula (see Methods and Figure S1 in S1 Text).

Since EFCM does not impose any constraints on fluxes, the enzyme-specific biomass production – and thus growth rate – is maximized by elementary flux modes, regardless of the values chosen for kinetic parameters [30, 31]. To see this, we consider all feasible steady-state flux modes, constrained to predefined flux directions and normalized to a unit biomass production rate. These flux modes form a convex polytope in flux space (see Figure 1(b)). The flux cost function is concave on this polytope [30], or even strictly concave for some rate laws [32], and so the minimal enzyme cost is achieved by a polytope vertex. In models without any active flux bounds, all these vertices are EFMs. Thus, to predict optimal flux modes, we need not scan all feasible flux modes, but can simply choose among EFMs. From our ECM calculations, we obtain the full spectrum of growth rates and yields of all EFMs. The rate/yield spectrum, a scatter plot between the two quantities, displays the possible trade-offs.

We now focus our attention on flux modes that maximize growth at a given yield, or maximize yield at a given growth rate. Such modes, which are not dominated by any other flux mode in terms of growth rate *and* yield, are called Pareto-optimal. They represent optimal compromises between growth rate and yield. If we could evaluate the growth rates and yields for *all* metabolic states in the model (including non-elementary flux modes), the resulting rate/yield points would form a dense, non-convex set. The border of this set, as drawn in Figure 1(a), is called the Pareto front. The EFMs on this front mark a selection of best compromises between growth rate and yield achievable in the model. By inspecting the rate/yield spectrum, we can tell whether there is an extended Pareto front or rather one metabolic state that optimizes both rate and yield. Even if growth and yield are positively correlated among all EFMs, the modes along the Pareto front will show a negative correlation whenever an extended front exists. Therefore, it is the size of the Pareto front that shows the extent of a rate/yield trade-off. While the yields are fixed properties of the EFMs, the growth rates depend on external conditions, and so does the rate/yield trade-off. We demonstrate this for a case study on *E. coli* bacteria, which have often been used for experiments on the rate/yield trade-off [9, 35–37] and whose enzyme kinetics are relatively well studied.

### Application of EFCM to *E. coli* core metabolism

To study growth rates and yields in *E. coli*, we applied EFCM to a model of core carbon metabolism. Our model, a modified version of the model presented in [38], comprises glycolysis, the Entner-Doudoroff pathway, the TCA cycle, the pentose phosphate pathway and by-product formation (see Figure 2(a), and Section 2 in S1 Text). The biosynthesis of macromolecules (“biomass”) from small metabolites and cofactors is not explicitly described, but summarized in an overall reaction for biomass production. Reaction kinetics are described by modular rate laws [39], and kinetic constants were obtained by parameter balancing [40] based on a large set of values reported in the literature (see Section 1.1 in S1 Text).

**Figure 2:**
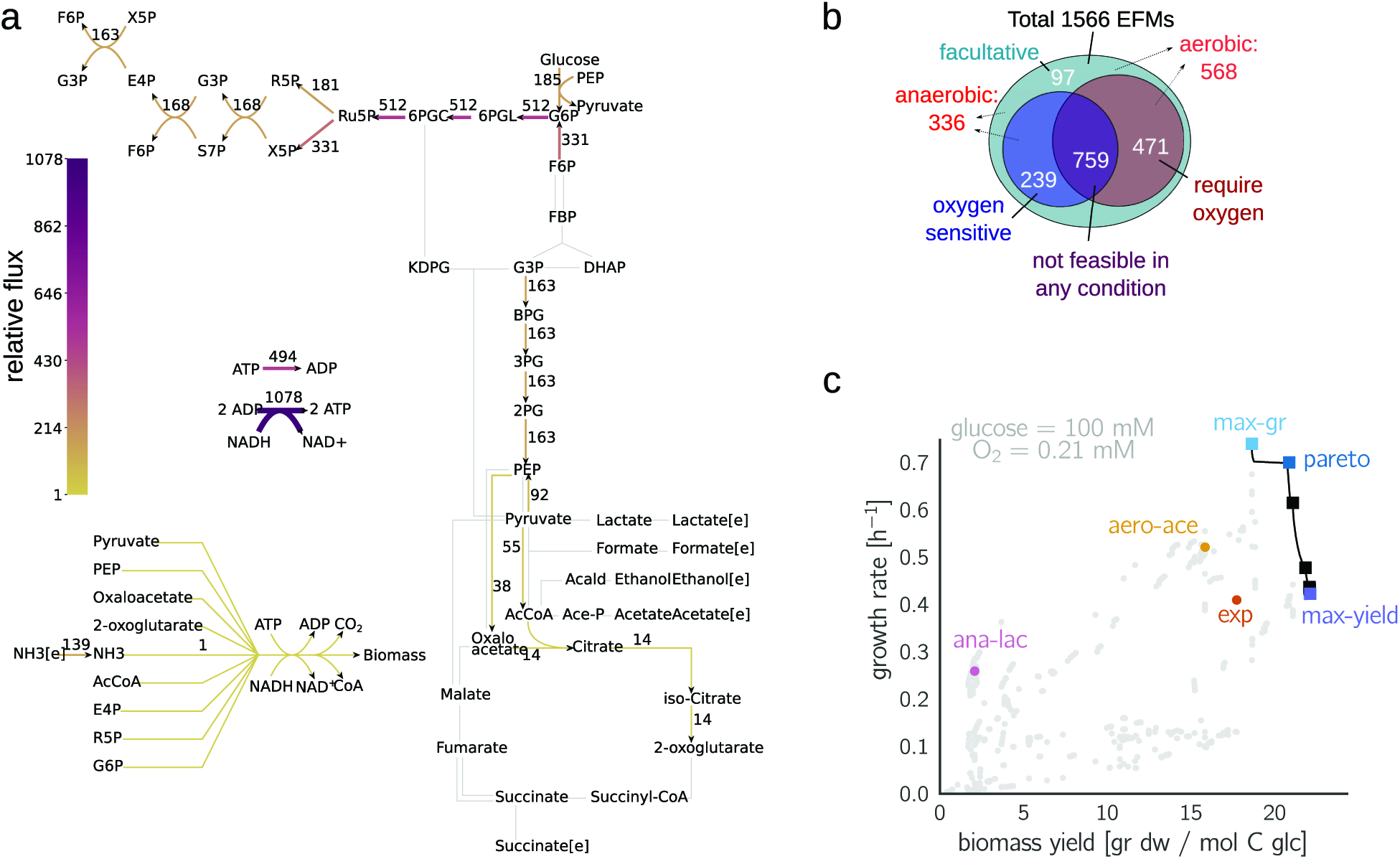
Metabolic strategies in *E. coli* metabolism. **(a)** Network model of core carbon metabolism in *E. coli*. Each Elementary Flux Mode (EFM) represents a steady metabolic flux mode in the network, scaled to a unit biomass flux. Reaction fluxes defined by the EFM *max-gr* are shown by colors. In our reference conditions – i.e. high extracellular glucose and oxygen concentrations – this EFM allows for the highest growth rate among all EFMs. Some of the cofactors in the model are not shown. **(b)** Statistics of biomass-producing EFMs. **(c)** Spectrum of growth rates and yields achieved by the EFMs. The labeled focal EFMs are described in Table 1, and their flux maps are given in Figures S25-S30 in S1 Text. Pareto-optimal EFMs are marked by squares; the Pareto front is shown by a black line. The plot reveals a positive correlation between growth rate and yield, despite the inevitably negative correlation among Pareto-optimal EFMs. See Figure S24 in S1 Text for a detailed view of the Pareto front and how it was sampled.

The yield of an EFM is defined as grams of biomass produced per mole of carbon atoms taken up in the form of glucose. EFMs that simultaneously use oxygen-sensitive enzymes (*pfl*) and oxygen-dependent reactions within the electron transport chain (*oxphos* or *sdh*) cannot be used by the cell. After discarding such EFMs, we obtained 568 EFMs that produce biomass under aerobic conditions and 336 under anaerobic conditions. 97 of these EFMs can operate under both conditions (Figure 2(b)). Statistical properties of the EFMs (size distribution, usage of individual reactions, and similarities between EFMs and measured fluxes) are shown in Figure S7 in S1 Text.

If all EFMs required the same total enzyme amount at *unit glucose uptake*, growth rates and yields would be proportional. Alternatively, if all EFMs required the same total enzyme amount at a *unit biomass production*, all EFMs would have exactly the same predicted growth rate, regardless of yield. Instead of these naïve approximations, we can now use our kinetic model and the EFCM method to obtain the actual spectrum of possible growth rates and yields (Figure 2(c)). While the yields are constant properties of the EFMs, the growth rates depend on enzyme demands and therefore on kinetics and extracellular nutrient levels. As reference conditions, we chose [glucose] = 100 mM, [O_2_] = 0.21 mM.

To visualize groups of similar EFMs, we used t-distributed Stochastic Neighbor Embedding (t-SNE), a machine learning algorithm for nonlinear dimensionality reduction [41]. The algorithm found five major clusters of EFMs, which loosely correspond to metabolic strategies (e.g. aerobic acetate-secreting EFMs). Since no kinetic information was used in t-SNE, we were surprised to find all EFMs with high growth rates in a single cluster (see Figure S6 in S1 Text).

To compare typical metabolic strategies, we focused on five EFMs with different characteristics and followed them across different external conditions and sets of kinetic parameters. We also show an experimentally determined flux distribution, called *exp* [42] (for calculations see Section 4.1 in S1 Text). These focal EFMs are marked by colors in Figure 2(b) and listed in Table 1. Flux maps (produced using software from [43]) can be found in Section 5.3 in S1 Text. The first three focal EFMs are located on the Pareto front. *max-yield*, the EFM with the highest yield, does not produce any by-products nor does it use the pentose-phosphate pathway. *max-gr* (whose flux map is shown in Figure 2(a)) has a slightly lower yield, but reaches the highest growth rate (0.739 h^−1^) in our reference conditions. It uses the pentose-phosphate pathway with a relatively high flux. In addition, we chose another EFM from the Pareto front (denoted *pareto*) with a growth rate and yield between the two extreme EFMs. Curiously, the EFMs along the Pareto front span only a narrow range of biomass yields (18.6 – 22.1), so there is almost no rate-yield trade-off. This is not a trivial finding, and other choices of parameters or extracellular conditions can lead to broader Pareto fronts: in low-oxygen conditions, the trade-off between growth rate and yield becomes much more pronounced.

**Table 1:** Focal EFMs representing different growth strategies. Metabolic fluxes are given in carbon moles (or O_2_ moles) per carbon moles of glucose uptake. Growths rate are given for reference conditions [glucose] = 100 mM, and [O_2_] = 0.21 mM. For more details, see Table S10 in S1 Text. Abbreviations: \**max-gr*: maximum growth rate; *max-yield*: maximum yield; *pareto*: a Pareto optimal EFM with higher growth rate than max-yield, and higher yield than max-gr; *ana-lac*: anaerobic lactate fermentation; *aero-ace*: aerobic acetate fermentation; *exp*: experimentally measured flux distribution.

| Acronym* | Biomass yield<br>(g/C-mol) | Growth rate<br>(h<sup>-1</sup>) | Oxygen<br>uptake | Acetate<br>secretion | Lactate<br>secretion |
| --- | --- | --- | --- | --- | --- |
| *max-gr* | 18.6 | 0.739 | 0.49 | 0 | 0 |
| *pareto* | 20.8 | 0.699 | 0.42 | 0 | 0 |
| *max-yield* | 22.1 | 0.422 | 0.39 | 0 | 0 |
| *ana-lac* | 2.1 | 0.258 | 0 | 0 | 0.92 |
| *aero-ace* | 15.8 | 0.520 | 0.21 | 0.35 | 0 |
| *exp* | 17.7 | 0.409 | 0.29 | 0.22 | 0 |

To study by-product formation, we consider two other EFMs below the Pareto front: an anaerobic lactate-fermenting mode (*ana-lac*) with a very low yield (2.1 g/C-mol) and an aerobic, acetate-fermenting mode (*aero-ace*) with a medium yield (15.2 g/C-mol). Interestingly, *ana-lac* has a ~10 times lower yield, but it still reaches about one third of the maximal growth rate, thanks to the lower enzyme cost of pentose phosphate pathway and lower glycolysis, as compared to TCA cycle and oxidative phosphorylation (per mol of ATP generated). This recapitulates a classic rate-versus-yield problem associated with overflow metabolism. Among all by-product forming EFMs, some acetate-producing EFMs have the highest growth rates, which might explain why *E. coli*, in reality, excretes acetate in aerobic conditions rather than lactate or succinate. Nevertheless, all by-product forming EFMs have lower growth rates than *max-gr* and are therefore not Pareto-optimal. Below we will see that this fact is subject to change when conditions are different, specifically at lower oxygen levels.

To study how by-product secretion affects yield and growth rate in general, we focused on some major uptake or secretion fluxes and visualized these fluxes for all EFMs in the rate/yield spectrum (Figure 3). EFMs close to the Pareto front consume intermediate amounts of oxygen and do not secrete any acetate, lactate or succinate. Another group of EFMs (shown in red in Figure 3(b)) consume slightly less oxygen, but secrete large amounts of acetate. Compared to pure respiration, these aerobic fermentation modes provides lower biomass yields. Other important fluxes are shown in Figure S8 in S1 Text.

**Figure 3:**
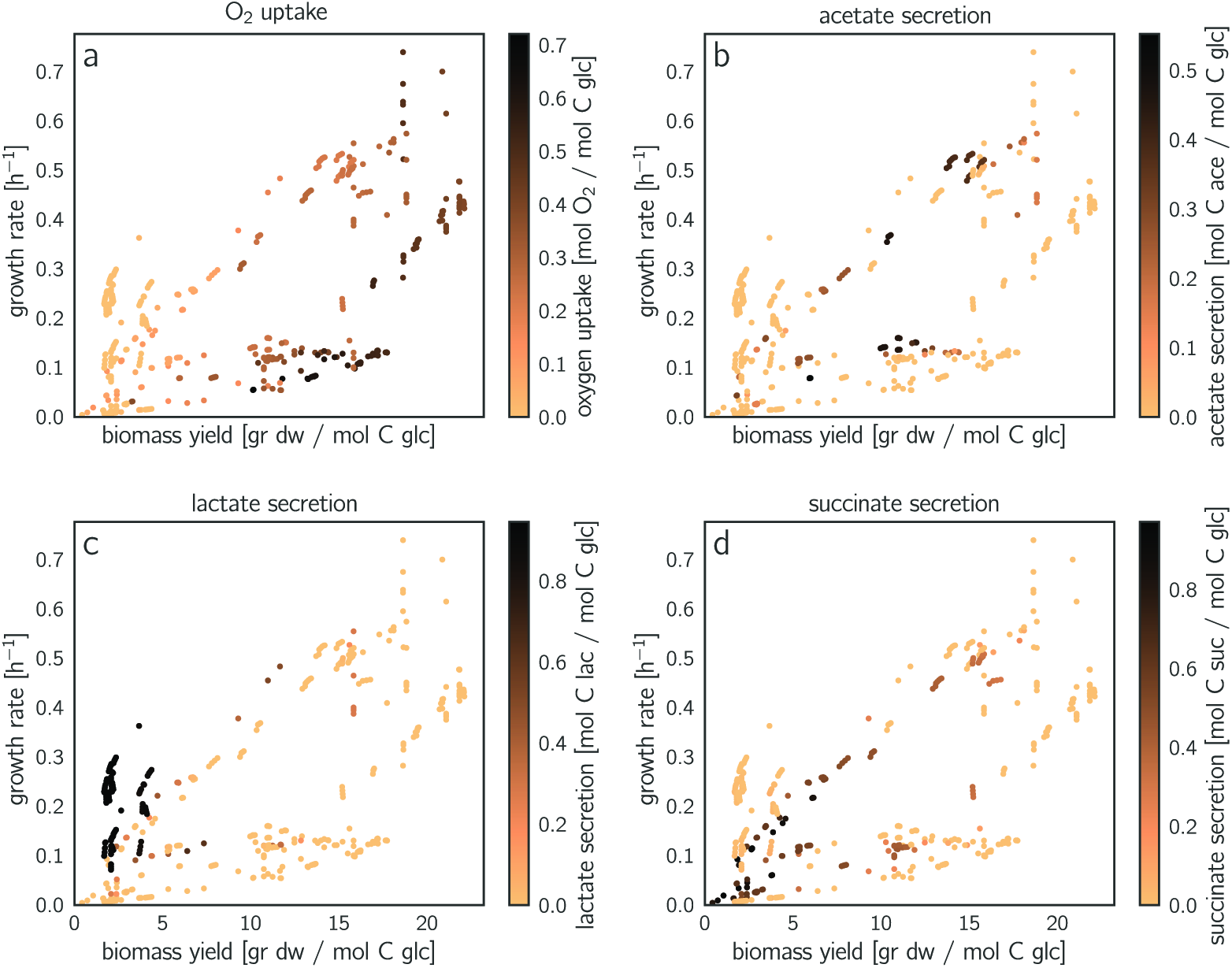
Uptake and secretion fluxes across EFMs. **(a)** Oxygen uptake (scaled by glucose uptake). Flux values are shown by colors in the rate/yield spectrum (same points as in Figure 2b). The EFMs with the highest growth rates consume intermediate levels of oxygen. The other diagrams show **(b)** acetate secretion, **(c)** lactate secretion and **(d)** succinate secretion, each scaled by glucose uptake. Acetate secretion and O_2_ uptake versus biomass yield are shown in Figure S9 in S1 Text.

### The effects of varying environmental conditions and varying enzyme parameters

The growth rate achieved by a flux mode depends on environmental conditions and enzyme parameters. To study this quantitatively, we varied some model parameters and traced their effects on the rate/yield spectrum. Figure 4(a) shows how lower oxygen levels affect the growth rate of oxygen-consuming EFMs. Lower oxygen levels need to be compensated by higher enzyme levels in oxidative phosphorylation, which lowers the growth rate (Figure 4(b) and Figure S16 in S1 Text). EFMs that function anaerobically, such as *ana-lac*, are not affected (see Figure S18 in S1 Text for enzyme allocation). Therefore, a low oxygen level leads to a prominent rate/yield tradeoff, with a Pareto front spanning a wide range of growth rates and yields (Figure 4(a)).

**Figure 4:**
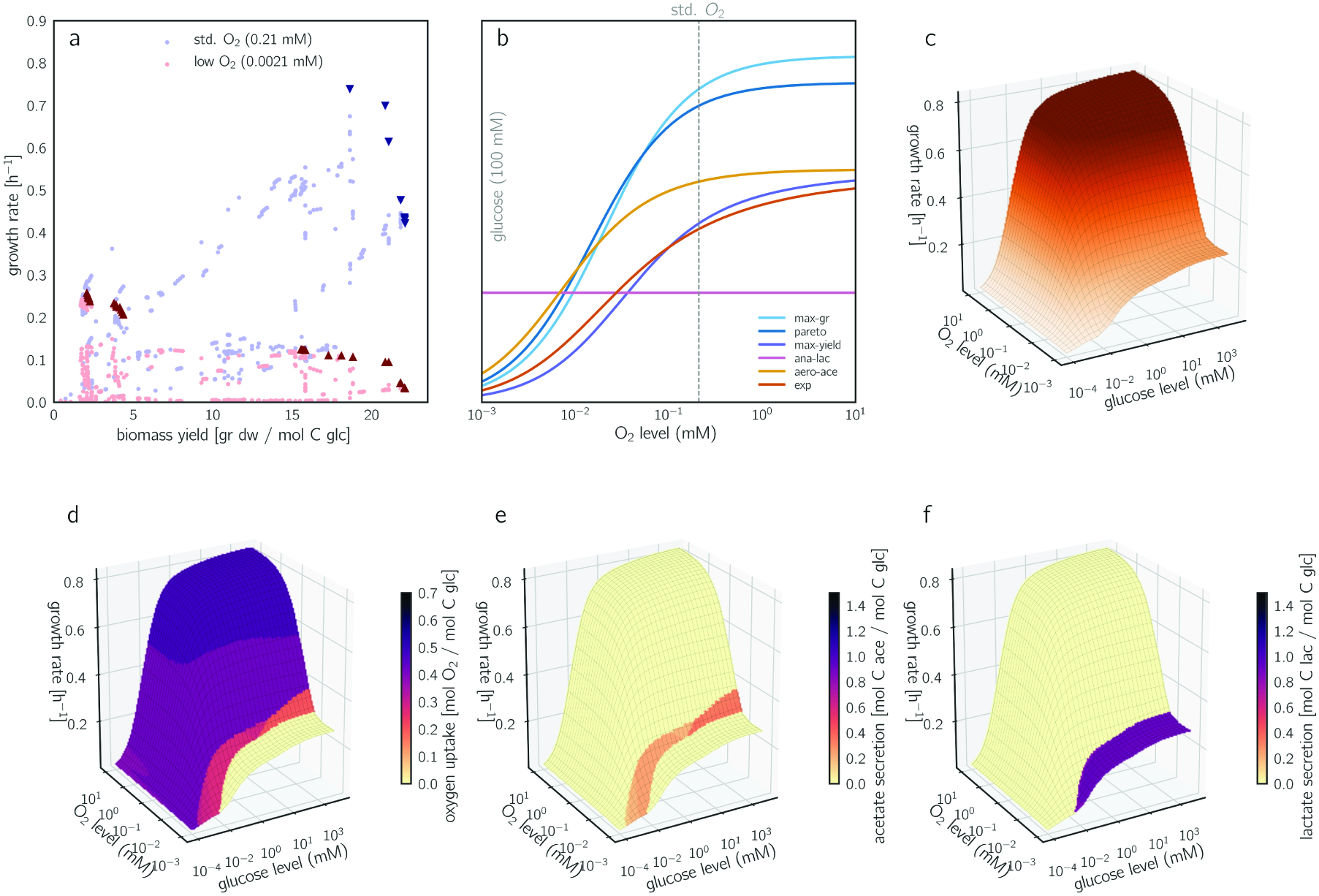
Growth rates and rate/yield trade-offs depending on glucose and oxygen levels. (a) Predicted growth rates and biomass yields of aerobic EFMs, at reference oxygen level (0.21 mM) and at a lower level (2.1 μM). Pareto-optimal EFMs are marked by dark triangles. Since changing oxygen levels affect the growth rate, but not the yield, points move vertically between the two conditions. Statistical distributions of growth rates across EFMs are shown in Figure S10 in S1 Text. (b) Oxygen-dependent growth rates for the five focal EFMs and the measured flux distribution. The oxygen level directly affects the catalytic rate of oxidative phosphorylation (reactions *oxphos* and *sdh*): lower oxygen levels require higher enzyme levels for compensation, to keep the fluxes unchanged. The non-respiring EFM *ana-lac* shows an oxygen-independent growth rate. In all other focal EFMs, the growth rate increases with the oxygen level and saturates around 10 mM. *max-gr*, which uses a higher amount of oxygen, has a steeper slope and loses its lead when oxygen levels drop below 1 mM. The corresponding changes in enzyme allocation are shown in Figure S18 in S1 Text. (c) Growth rate as a function of glucose and oxygen levels (“Monod surface”). For a closed approximation formula, see Section 4.6 in S1 Text. (d)-(f) The same plot, with oxygen uptake, acetate secretion, and lactate secretion shown by colors. Distinct areas represent different optimal EFMs (compare Figure S13 in S1 Text). The optimal EFMs for strictly anaerobic conditions are depicted in Figure S15 in S1 Text (b).

The effect of external glucose levels can be studied similarly (Figures S12 and S16 in S1 Text): at lower external glucose concentrations, the PTS transporter becomes less efficient and cells must increase its expression in order to maintain the flux. This increases the total enzyme cost and slows down growth. Below a glucose concentration of 10^−3^ mM, the demand for transporter dominates the enzyme demand completely (see Figure 5(b) and Figures S17-S18 in S1 Text for a breakdown of enzyme allocation). Since the PTS transporter is the only glucose transporter in our model, it is used by all EFMs, leading to a universal monotonic relationship between glucose concentration and growth rate. However, the detailed shape of the glucose/growth rate plot, known as the Monod curve [44, 45], depends on the PTS flux and on many other parameters that differ between EFMs (see Section 3.3 in S1 Text)). The performance of EFMs under high-glucose and low-glucose conditions is shown in Figure S19 in S1 Text.

**Figure 5:**
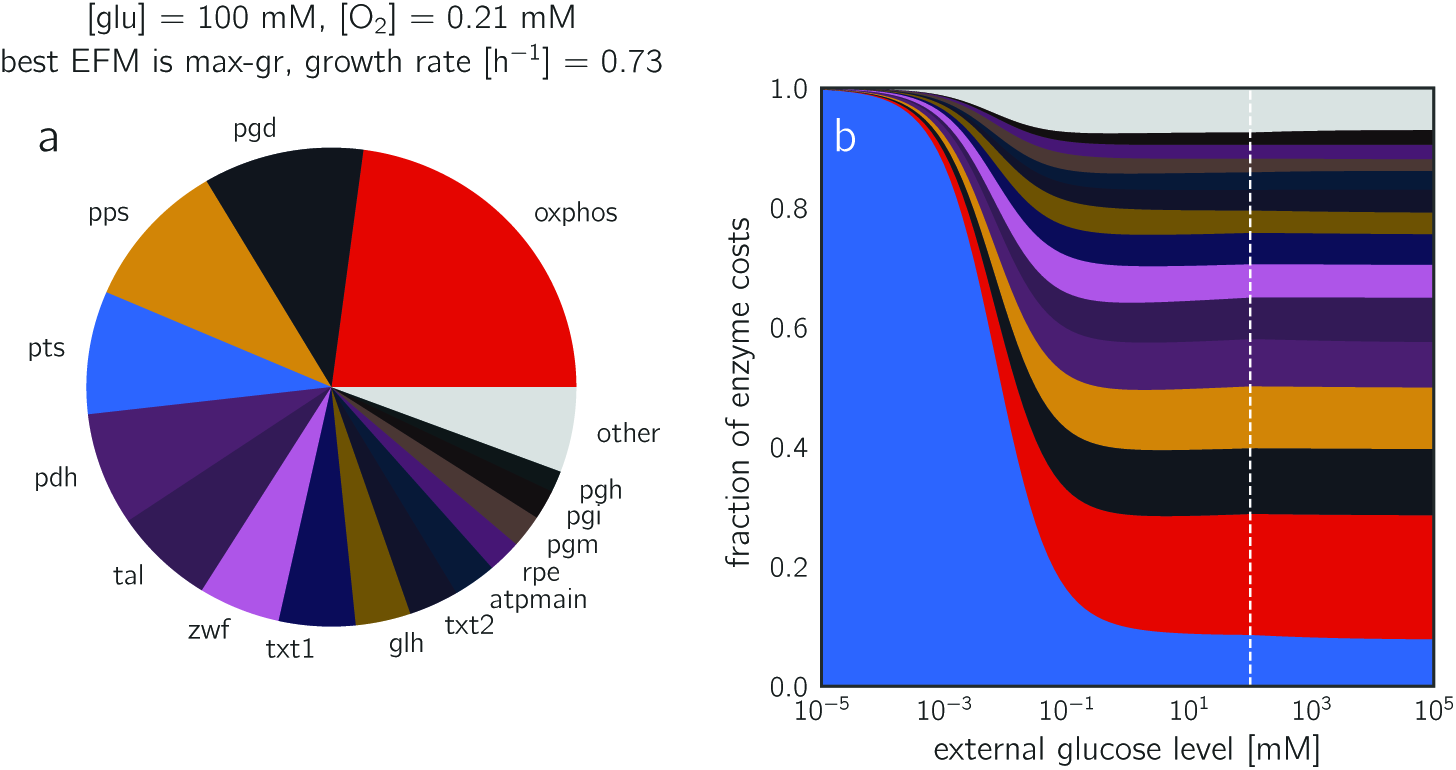
Predicted protein investments. (a) Predicted protein demands for the EFM *max-gr* at reference conditions. (b) Predicted protein demand for the EFM *max-gr* at varying glucose levels and reference oxygen level. The y-axis shows relative protein demands (normalized to a sum of 1). The dashed line indicates the reference glucose level (100 mM) corresponding to the pie chart in panel (a).

By varying the glucose and oxygen levels, we can screen a range of environmental conditions and obtain a two-dimensional Monod surface plot. The winning strategies, i.e. the EFMs with the highest growth rates can be depicted on this surface (Figure 4 (d–e)) or in a glucose/oxygen phase diagram (see Figures S13-S15 in S1 Text, also for anaerobic conditions). More than 20 different EFMs achieve a maximal growth rate in at least one of the conditions scanned. To simplify the picture, we can focus on EFM features such as uptake rates and plot them on the Monod surface (Fig 4(c)-(f)). As expected, oxygen uptake (Figure 4(d)) decreases when oxygen levels are low. This pattern occurs across the entire range of glucose levels, but the transition – from full respiration to acetate overflow (Figure 4(e)) and further to anaerobic lactate fermentation EFMs (Figure 4(f)) – is shifted at lower glucose levels. Interestingly, this transition disappears at extremely low glucose concentrations (0.1 μM), as the fully respiring *pareto* EFM exhibits the highest growth rate even at the lowest oxygen levels tested (Figure S13(a) in S1 Text).

While glucose levels are relatively easy to adjust in experiments, it is difficult to measure oxygen levels in the local environment of exponentially growing cells. This has resulted in a long-standing debate about the exact conditions that *E. coli* cells experience in batch cultures [46–48], and it makes it hard to validate our predicted transition from acetate fermentation to full respiration. Our model predicts that at a constant level of [O_2_], *E. coli* will fully respire at low glucose levels and secrete acetate at high glucose levels (see Figure 4). A similar shift from pure respiration to a mixture of respiration and acetate secretion has been observed in chemostat cultures [49], where higher glucose levels result from higher dilution rates.

The choice of metabolic strategies does not only depend on external conditions, but also on enzyme parameters. As an example, we varied the *k*_*cat*_ value of triose-phosphate isomerase (*tpi*) and traced changes in the rate/yield spectrum. Not surprisingly, slowing down the enzyme decreases the growth rate (see Figure S20 in S1 Text). But to what extent? Two of our focal EFMs (*max-gr* and *pareto*) are not affected at all, since they do not use the *tpi* reaction. All other focal EFMs show strongly reduced growth rates. To study this systematically, we predicted the growth effects of *all* enzyme parameters in the model (equilibrium constants, catalytic constants, Michaelis-Menten constants) by computing the growth sensitivities, i.e. the first derivatives of the growth rate with respect to the enzyme parameter in question (see Section 4.2 in S1 Text, and supplementary data files). A sensitivity analysis between all model parameters and the growth rates of all EFMs (or alternatively, their biomass-specific enzyme cost) can be performed without running any additional optimizations (Sections 4.3 – 4.4 in S1 Text). Growth sensitivities are informative for several reasons. On the one hand, parameters with a large impact on growth will be under strong selection (where positive or negative sensitivities indicate a selection for larger or smaller parameter values, respectively). On the other hand, these are also the parameters that need to be known precisely for reliable growth predictions. The parameters of a reaction can have very different effects on the growth rate. For example, the sensitivities of the *k*_*cat*_ and *K*_*M*_ values of *pgi* are low, but the growth rate is very sensitive to the *K*_*eq*_ value.

To study the effects of a gene deletion, we can simply discard all EFMs that use the affected reaction: based on a precalculated EFCM analysis of the full network, we can easily analyze the restricted network without any new optimization runs. By switching off pathways, we can easily quantify the growth advantage they convey. Instead of studying pathways in isolation as in *Flamholz et al.* [21], we can study their usage as part of a whole-network metabolic strategy. Figure 6 shows an analysis for two common variants of glycolysis, the (high ATP yield, high enzyme demand) EMP and the (low ATP yield, low enzyme demand) ED pathway, across different external glucose and oxygen levels (see Section 3.4 in S1 Text). At low oxygen levels and medium-high glucose levels (10 μM – 100 mM), cells profit strongly from using the ED pathway, and knocking it out decreases the growth rate by up to 25%. The EMP pathway provides a much smaller advantage (up to 10%), and only in a narrow range of low-oxygen conditions.

**Figure 6:**
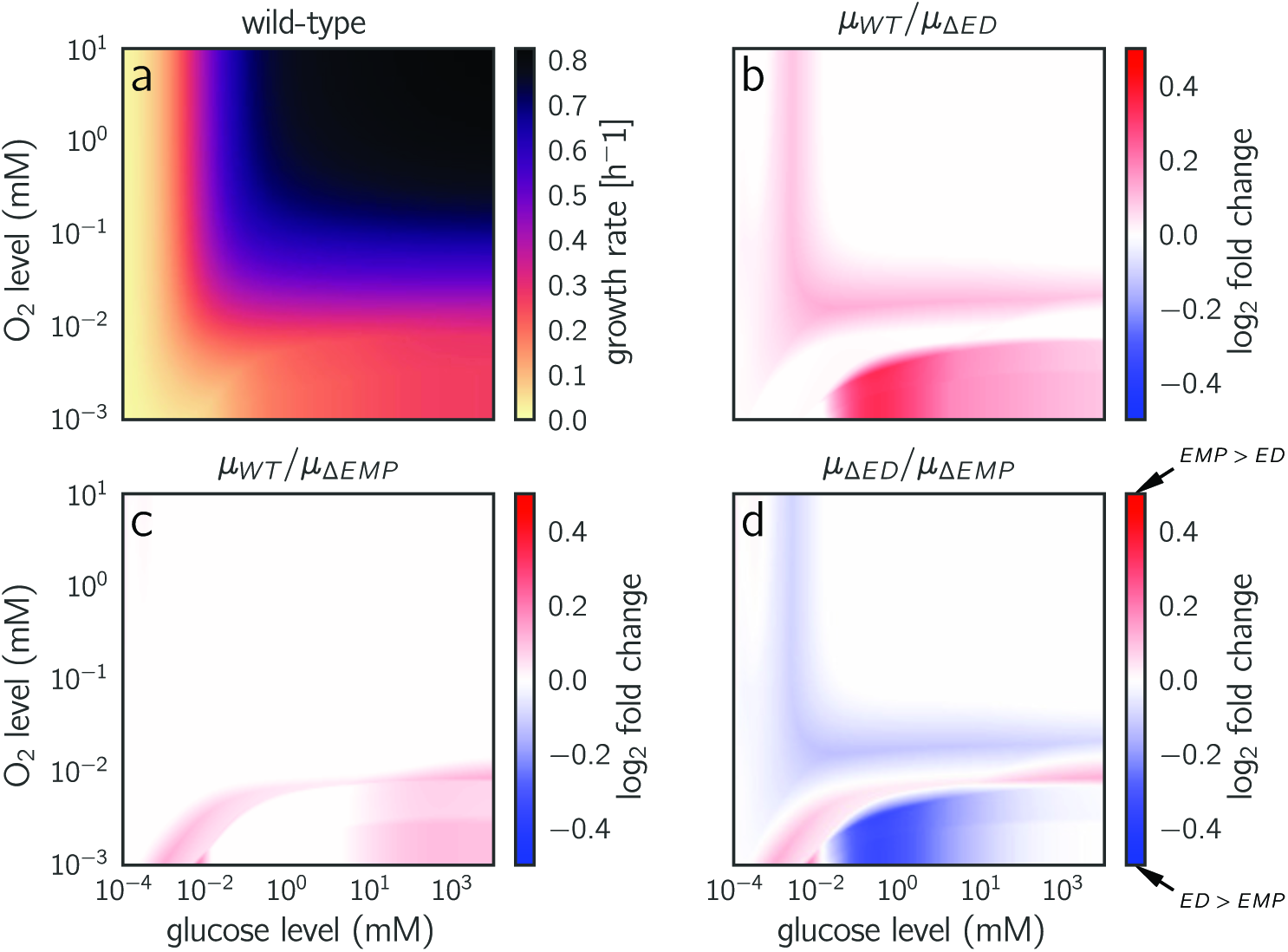
Growth rates achieved with two variants of glycolysis. (a) Glucose- and oxygen-dependent growth rates predicted for wild-type *E. coli*. Same data as in Figure 4(c), but shown as a heatmap. *E. coli* can employ two variants of glycolysis: the Embden-Meyerhof-Parnas (EMP) pathway, which is common also to eukaryotes, and the Entner-Doudoroff (ED) pathway, which provides a lower ATP yield at a much lower enzyme demand [21]. (b) A simulated ED knockout strain that must use the EMP pathway. The heatmap shows the relative growth advantage of the wild-type strain (i.e. of reintroducing the ED pathway to the cell). The ED pathway provides its highest advantage at low oxygen and medium to low glucose levels. (c) Growth advantage provided by the EMP pathway. The advantage is highest at glucose concentrations below 10 μM. (d) Comparison between the two knockout strains. Blue areas indicate conditions where ED is more favorable, and red areas indicate conditions where EMP would be favored. The dark blue region at low oxygen and medium glucose levels may correspond to the environment of bacteria such as *Z. mobilis*, which uses the ED pathway exclusively [50]. The same data are shown as Monod surface plots in Figure S21 in S1 Text.

## Discussion

Our case study on *E. coli* metabolism reinforces the notion that growth rate and biomass yield are not strictly coupled. Instead, their correlations across EFMs, and the extent of rate/yield trade-offs along the Pareto front, depend on details such as growth conditions and enzyme parameters. At high oxygen levels, growth-maximizing flux modes have an almost maximal yield and the Pareto front is very narrow. In contrast, under low-oxygen conditions the highest growth rates are obtained by low-yield strategies and a long Pareto front emerges (Figure 4(a)). It is not surprising that experimental results indicating rate/yield trade-offs were inconclusive and difficult to interpret. As shown in [9], wild-type cell populations might be far from the Pareto front, and a selection for fast growth may push the populations and individuals closer to it. It would be interesting to study whether these results are in fact dependent on oxygen availability.

EFCM predicts which flux modes are likely to be used by well-adapted cells. We expected that the EFM with the highest growth rate (*max-gr*, in the standard conditions chosen in this study) would coincide with the experimentally determined flux mode (*exp*) in the same conditions. However, this is not the case, and the two flux modes are not even very similar (correlation *r* = 0.41, see Figure S7(c) in S1 Text). Our model predicts a much higher maximal biomass yield than the yield measured in batch cultures (18.6 vs 11.8 gr dry weight per carbon mole [51]), while the predicted growth rate is slightly lower (0.74 vs 0.89 h^−1^). However, for the experimentally determined flux mode (*exp*), we overestimate the yield (17.7 vs 11.8 [42]) and underestimate the growth rate (0.41 vs 0.89) as well, so some of the discrepancies may be due to weaknesses of our model (e.g. wrong kinetic parameter values) rather than due to EFCM itself. The overestimation of yield (which depends on network structure, not on kinetics) may be caused by the fact that our model misses some waste products or additional processes that dissipate energy, or that our high-yield EFMs are kinetically unfavorable in reality. The underestimated growth rates may result from our simplistic conversion of enzyme costs into growth rates. However, we hope that these over-and underestimations occur consistently across EFMs and do not affect the qualitative results of this study.

In contrast to the much simpler model by Basan *et al.* [49], our model does not predict growth-rate dependent acetate overflow as observed in *E. coli*. In our standard aerobic conditions (see Figure 2 and Figure S14(h)) in S1 Text, the winning mode, *max-gr*, is completely respiratory and produces no fermentation products. Only at low oxygen levels, EFMs with acetate overflow, such as *aero-ace*, become favorable (see Figure 4 and Figure S15(e) in S1 Text). This misprediction may depend on several factors:

First, we may have underestimated the effective cost of oxidative phosphorylation (*oxphos*), which becomes costly at lower oxygen levels, or we may have overestimated the oxygen availability. The oxygen concentration of [O_2_] = 0.21 mM, which we chose to represent typical laboratory conditions, may be inaccurate; oxygen availability may be as complex as in yeast, where it seems to diffuse too slowly to supply the mitochondria fully with oxygen [48]. Moreover, the affinity of the reactions to oxygen is not precisley known, so even a precise value of the oxygen concentration would not suffice.

Second, the experimentally observed acetate production may result from additional, growth-rate dependent flux constraints like those employed by Basan *et al.* in their model. In our model, we did not impose any bounds on fluxes (aside from normalizing the flux modes to unit per biomass production), and thus metabolic efficiency is maximized by an EFM. The growth rate does not even appear in the optimization. We account for it only later, when metabolic efficiency is translated into an achievable growth rate. Thus, it is possible that we miss some physiological constraints such as membrane real-estate [52], changing biomass composition, or extracellular oxygen diffusion rates. Even without flux constraints, some EFMs mix respiration and acetate production, e.g. *aero-ace*. However, none of them corresponds exactly to the fluxes observed experimentally. Moreover, the measured relative rate of acetate production increases continuously with the growth rate, which cannot be captured by a single constant EFM. A usage of flux constraints in EFCM would be possible and would allow us, for example, to limit certain fluxes or to enforce some minimal flux, e.g. in ATP-consuming maintenance reactions. To screen all vertices of the flux polytope, one may build on the concept of elementary flux vectors [53, 54]. However, the number of these vertices may become very large, and whenever flux bounds are changing (e.g. as a function of growth rate), this would change the set of polytope vertices, and the entire calculation would have to be done for each growth rate.

Third, it is also possible that the experimentally observed acetate secretion is simply not optimal. In adaptive laboratory evolution experiments [36, 37], the evolved strains grew about 1.5 times faster without a significant change in yield, but most of this increase could be explained by an increasing glucose uptake because the relative rates of acetate overflow did not change. Apparently, if acetate secretion is due to a glucose uptake constraint, this constraint can be bypassed by mutations and cells may be able to decrease acetate secretion while growing faster. In a recent comparison of seven *E. coli* wild-type strains [35], three strains were found to secrete no acetate at all in aerobic conditions (on glucose), but to use a fully respiratory strategy without any by-product secretion. Two of these fully respiring strains grew just as fast as the evolved strains from the adaptive evolution studies (about 1.0/h), and significantly faster than the lab strain that we used for our reference flux data and for the stoichiometric model (K-12). Again, this finding raises questions about universal rate/yield trade-offs and supports our conclusion that the trade-off may almost disappear in high-oxygen conditions.

Some variants of FBA manage to predict flux distributions with a suboptimal biomass yield by putting bounds on enzyme investments. An example is FBAwMC (Flux Balance Analysis with Molecular Crowding), which relates fluxes to enzyme demands and limits the cytoplasmatic protein density [55]. However, these methods are insensitive to environmental conditions: the crowding coefficients assigned to reactions are constants, and metabolite concentrations are not considered at all. In [20], Müller et al. ran a kinetic optimization (which attempts to solve the nonlinear enzyme minimization problem directly) and compared it to a linear approximation called satFBA. In this approximation, the constraints are exactly like in FBAwMC, except that the crowding coefficients of exchange reactions are divided by saturation values. The saturation values, numbers between 0 and 1, account for the concentrations of external metabolites such as glucose and oxygen. For a small metabolic network (comprising 5 reactions), satFBA yields the same qualitative predictions as a kinetic optimization (and EFCM, for that matter), in particular with regard to the rate/yield trade-off. However, satFBA assumes that transport reactions are the only reactions affected by metabolite levels, whereas EFCM models the interplay between metabolite levels, enzyme efficiencies, and enzyme investments in all enzymatic reactions. It remains to be seen whether satFBA, with its single kinetic bottleneck, can reproduce complex predictions of EFCM like the ones shown in Figure 4.

Constraint-based whole-cell models such as Resource Balance Analysis (RBA) [56, 57] or ME-models [58] treat protein production as a part of the cellular network and couple metabolic rates to production rates of the catalyzing enzymes. These methods differ from EFCM in three main ways: in the modeling of protein production, of catalytic rates, and of biomass composition and enzyme cost weights. (i) While RBA and ME model protein production in detail, EFCM is limited to metabolism: the partitioning between metabolic enzymes and ribosomes is captured by a formula that effectively converts enzyme cost into growth rate (see Methods). (ii) In reality, enzymes often operate below their maximal speed (i.e. the *k*_cat_ value), at a catalytic rate called apparent *k*_cat_ value [59]. This capacity utilization lower than 1 depends on metabolite levels and is quantified by the efficiency factors of ECM [34]. For each enzyme, the capacity utilization computed by EFCM varies across EFMs, but remains close to some typical value. These values, for different enzymes, span almost the entire range between 0 and 1 (see Figure S11 in S1 Text). In a linearized variant of EFCM that assumes full capacity utilization, the growth rate would be overestimated and the growth differences between EFMs would be distorted. In fact, our predicted enzyme cost is between 1.4 and 4.7 times higher (depending on the EFM considered) than the ideal costs of enzymes operating at their maximal capacity (see Figure S3 in S1 Text). RBA avoids this problem by replacing the *k*_cat_ values by empirically determined, growth-rate dependent apparent catalytic rates. Constraint-based methods that ignore this effect [23, 60] underestimate the actual enzyme demand, thus suggesting an “unused enzyme fraction” in cells [61]. We think that “unexplained enzyme fraction” would be a better term, because the enzyme amount predicted for fully efficient enzymes is an ideal value that would simply not suffice to catalyze the required fluxes in reality, given all thermodynamic and kinetic constraints [34, 62]. (iii) In contrast to RBA and ME models, EFCM assumes a fixed biomass composition and fixed cost weights for the enzyme molecules. This means that cells, in EFCM, lack some strategic options that exist in RBA and ME models: to fine-tune the biomass composition towards a usage of “cheap” precursors, or to decrease the cost weights of proteins by cost-optimizing the production of limiting protein components such as iron. Again, these options would be hard to implement in EFCM because biomass composition is a defining part of the stoichiometric model, and any growth-rate dependent changes in biomass composition would also change the set of EFMs.

Although efficient protein allocation may be important for fast growth [63], there is empirical evidence that cells do not always minimize enzyme cost. *Lactococcus lactis*, for example, can undergo a metabolic switch that leads to big changes in growth rate, but involves no changes in protein levels [64]. These cells could, in theory, save enzyme resources while maintaining the same metabolic fluxes, but do not do so – possibly because their enzyme levels provide other benefits, e.g. anticipating metabolic changes to come. EFCM ignores such complex objectives: it describes fully optimal, but “short-sighted” cell strategies which define a lower bound on the enzyme demand. By considering secondary objectives, e.g., a need for preemptive protein expression or safety margins to counter expression fluctuations, one would predict higher demands and lower growth rates.

Our study has demonstrated that enzyme kinetics is a useful addition to constraint-based flux prediction (see Section 1.4 in S1 Text)). In contrast to the minimal model in [49], our model was not fitted to recapitulate a specific known phenomenon, but was made to derive predictions *ab initio* in the spirit of “testing biochemistry” [65]. As long as *in vivo* kinetic constants are not precisely known, this harbours the risk of mispredictions. Curiously, for example, the EFMs with the highest predicted growth rates bypass upper glycolysis and use the pentose phosphate pathway instead. On the contrary, an *ab initio* approach allows modelers to recover empirical laws directly from cell biological knowledge, for example, the shape of Monod curves and Monod surfaces (see Figure S15 and Section 4.6 in S1 Text for general simplified Monod functions). It allows us to compute quantitative effects of allosteric regulation or mutated enzymes (see Figure S2 in S1 Text), the residual glucose concentration in chemostats (see Figure S15 in S1 Text), and the trade-offs between metabolic strategies at different glucose levels (see Figure S19 in S1 Text). The decomposition into EFMs also greatly facilitates calculating the epistatic interactions between reaction knockouts (see Figure S2 (f) in S1 Text). Although yield-related epistatic interactions were previously computed using FBA (see Section 3.5 in S1 Text), environment-dependent epistatic effects on growth rate have not been computed so far. EFCM could be applied to larger models and models with flux constraints, and other cost functions could be implemented (see Section 1.6 in S1 Text). As a fully mechanistic method, it puts existing biochemical models and ideas about resource allocation to test and enables us to address fundamental issues of unicellular growth and cell metabolism, such as the trade-off between growth rate and biomass yield.

## Methods

### Optimal enzyme and metabolite profiles

A metabolic state is characterized by cellular enzyme levels, metabolite levels, and fluxes. All these variables are coupled by rate laws, which depend on external conditions and enzyme kinetics. The EFCM algorithm finds optimal metabolic states in the following way. First, we enumerate the elementary flux modes of a network, which constitute the set of potentially growth-optimal flux modes. Then we consider a specific simulation scenario, defined by kinetic constants and external metabolite levels, and compute the growth rates for all EFMs. To determine the optimal metabolic state – a state expected to evolve in a selection for fast growth – we choose the EFM with the highest growth rate.

The optimal state (**v, c, E**) can be found efficiently by a nested screening procedure (Figure 1(b-c)). First, we consider all EFMs, normalized to a given biomass production rate *v*_BM_. To determine the relative enzyme demand of an EFM, we predefine *v*_BM_, scale our EFM to realize this production rate, and compute the enzyme demand by applying Enzyme Cost Minimization (ECM), i.e. an optimization of metabolite levels c and enzyme levels E. ECM has recently been applied to a similar model of *E. coli*’s core carbon metabolism [34]. It assumes a given flux distribution (in our case, an EFM) and treats the enzyme concentrations as explicit functions of substrate and product levels and fluxes. Given a flux mode v, we consider all feasible possible metabolite profiles lnc, consistent with the flux directions and respecting predefined bounds on metabolite levels. For each such profile, we compute the enzyme demands *E*_*l*_ and the total enzyme mass concentration *E*_met_ = Σ_*i*_ *ω*_*i*_ *E*_*i*_(in mg l^−1^), where *ω*_*i*_ denotes the molecular mass of enzyme *l* in Daltons (mg mmol^−1^) and enzyme concentrations are measured in mM (i.e., mmol l^−1^). As a function of the logarithmic metabolite levels, *E_met_* is convex; this allows us to find the global minimum efficiently. In the model, we use common modular rate laws [39], for which the enzymatic cost in log-metabolite space is strictly convex (Joost Hulshof, personal communication). The optimized enzyme cost is a concave function in flux space [30–32]. This combination of convexity and concavity allows for a fast optimization of enzyme levels and fluxes for each condition and set of kinetic parameters.

### Online tool for Enzyme Cost Minimization

We implemented ECM in the Network-Enabled Optimization System (NEOS), an internet-based client-server application that provides access to a library of optimization solvers. The NEOS Server is available free of charge and offers a variety of interfaces for accessing the solvers, which run on distributed high-performance machines enabled by the HTCondor software. The NEOS Guide website (https://neos-guide.org) showcases optimization case studies, presents optimization information and resources, and provides background information on the NEOS Server. Using our online service, users can run EFCM for their own models, using different rate laws. With our *E. coli* model, the optimization for one flux distribution takes a few seconds, and for the complete set of all EFMs several minutes on a shared Dell PowerEdge R430 server with 32 intel xeon cores. Details can be found in Section 1.2 in S1 Text, and on the web page (www.neos-guide.org/content/enzyme-cost-minimization).

### Converting enzyme-specific biomass rates into growth rates

Following the approach of Scott et al. [66], cell growth rates can be predicted from the demand for metabolic enzyme, divided by the rate of biomass production (see Section 1.3 in S1 Text)). A cell’s growth rate is given by *μ* = *v*_BM_/*c*_BM_, where *c*_BM_ is the biomass amount per cell volume and *v*_BM_ is the biomass production rate (biomass amount produced per cell volume and time). If cell biomass consisted only of metabolic enzymes (more precisely, of enzymes considered in the cost *E*_met_), the enzyme-specific biomass production rate *r*_BM_ = *v*_BM_/*E*_met_ would be equal to the cellular growth rate. Since this is not the case, we convert between *E*_met_ and *c*_BM_ using the approximation*E*_met_/*c*_BM_ = *ƒ*_prot_(*a − b μ*), where *ƒ*_prot_ = 0.5 is the fraction of protein mass within the cell dry mass and the parameters *a* = 0.27 and *b* = 0.2h were fitted to describe the metabolic enzyme fraction in proteomics data, assuming a linear dependence on growth rate [66]. As shown in the S1 Text (Equations 8-9 and Figure S1), we obtain the conversion formula

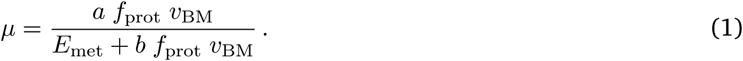

Note that the biomass flux *v*_*R70*_ in our model is set to 1 mM s^−1^ by convention, and the *k*_*cat*_ of this reaction was set to a sufficiently high value so that it would never become a bottleneck (see Figure S5 in S1 Text). By simple unit conversion we obtain *v*_BM_ = 7.45 x 10^7^ mg l^−1^ h^−1^. As shown above, the total enzyme mass concentration is given by *E*_met_ = Σ_*i*_ *ω*_*i*_ *E*_*i*_ in units of mg l^−1^, so it requires no further conversion. The final formula for growth rates, with proper units, reads

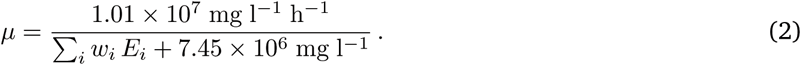

It shows that maximizing the growth rate *μ* is equivalent to minimizing the enzyme cost *E*_met_. The link between biomass production, total enzyme mass concentration, and growth rate can also be understood through the cell doubling time. We first define the enzyme doubling time 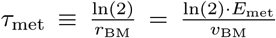 the doubling time of a hypothetical cell consisting only of core metabolism enzymes. Since *E. coli* cells contain also other biomass components, the real doubling time is longer and depends on the fraction of these other components within the total biomass. Furthermore, this fraction decreases with the doubling time, as seen in experiments [67] and as expected from trade-offs between metabolic enzymes and ribosome investment [66]. This leads to a constant offset in the final cell doubling time formula:

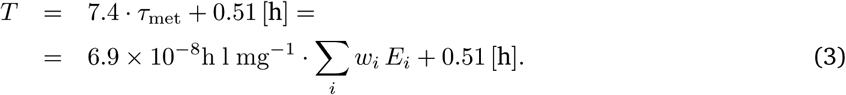

### Growth rate sensitivities

The calculation of sensitivities between enzyme parameters and growth rate is based on the following reasoning. If a parameter change slows down a reaction rate, this change can be compensated by increasing the enzyme level in the same reaction while keeping all metabolite levels and fluxes unchanged. For example, when a catalytic constant changes by a factor of 0.5, the enzyme level needs to be increased by a factor of 2. The cost increase is given by Δ cost 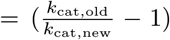 ·[old enzyme cost]. Also for other parameters, the local enzyme increase can be simply computed from the reaction’s rate law. Instead of adapting only one enzyme, the cell may save some costs by adjusting all enzyme and metabolite levels in a coordinated fashion. However, the extra cost advantage is only a second-order effect and can be neglected for small parameter variations. Hence, the first-order local and global cost sensitivities are completely identical (proof in Section 4.2 in S1 Text). Sensitivities to external parameters (e.g. extracellular glucose concentration) can be computed similarly. The growth sensitivities for a given EFM are computed by multiplying the enzyme cost sensitivities by the derivative between growth rate and enzyme cost.

### Supporting Information

**S1 Text Supplementary text containing Figures S1 - S30, Tables T1 - T10, and a list of supplementary data files available on GitHub.**

## Acknowledgments

We thank H.-G. Holzhütter, Joost Hulshof, Daan de Groot, Avi Flamholz, Philip van Kuiken, Timo Maarleveld and Bas Teusink for fruitful discussions, and the Lorenz Center of Leiden University for providing a space for developing ideas.

## References

[1] Goel A, Wortel MT, Molenaar D, Teusink B. Metabolic shifts: A fitness perspective for microbial cell factories. Biotechnology letters. 2012;34(12):2147–2160.

[2] Pfeiffer T, Schuster S, Bonhoeffer S. Cooperation and competition in the evolution of ATP-producing pathways. Science. 2001;292(5516):504–507.

[3] MacLean RC. The tragedy of the commons in microbial populations: Insights from theoretical, comparative and experimental studies. Heredity (Edinb). 2008;100(5):471–477.

[4] Werner S, Diekert G, Schuster S. Revisiting the thermodynamic theory of optimal ATP stoichiometries by analysis of various ATP-producing metabolic pathways. Journal of Molecular Evolution. 2010;71(5-6):346–355.

[5] Westerhoff HV, Hellingwerf KJ, van Dam K. Thermodynamic efficiency of microbial growth is low but optimal for maximal growth rate. Proceedings of the National Academy of Sciences. 1983;80(1):305–309.

[6] Molenaar D, van Berlo R, de Ridder D, Teusink B. Shifts in growth strategies reflect tradeoffs in cellular economics. Molecular Systems Biology. 2009;5(1):323.

[7] Noor E, Bar-Even A, Flamholz A, Reznik E, Liebermeister W, Milo R. Pathway thermodynamics highlights kinetic obstacles in central metabolism. PLoS Computational Biology. 2014;10.

[8] Velicer GJ, Lenski RE. Evolutionary trade-offs under conditions of resource abundance and scarcity: Experiments with bacteria. Ecology. 1999;80(4):1168–1179.

[9] Novak M, Pfeiffer T, Lenski RE, Sauer U, Bonhoeffer S. Experimental tests for an evolutionary trade-off between growth rate and yield in E. coli. The American Naturalist. 2006;168(2):242–251.

[10] Prasad RM, Seeto S, Ferenci T. Divergence and redundancy of transport and metabolic rate-yield strategies in a single Escherichia coli population. Journal of Bacteriology. 2007;189(6):2350–2358.

[11] Fitzsimmons JM, Schoustra SE, Kerr JT, Kassen R. Population consequences of mutational events: Effects of antibiotic resistance on the r/K trade-off. Evolutionary Ecology. 2010;24(1):227–236.

[12] Bachmann H, Fischlechner M, Rabbers I, Barfa N, dos Santos FB, Molenaar D, et al. Availability of public goods shapes the evolution of competing metabolic strategies. Proceedings of the National Academy of Sciences. 2013;110(35):14302–14307.

[13] Brown GC. Total cell protein concentration as an evolutionary constraint on the metabolic control distribution in cells. Journal of Theoretical Biology. 1991;153(2):195–203.

[14] Tepper N, Noor E, Amador-Noguez D, Haraldsdóttir HS, Milo R, Rabinowitz J, et al. Steady-State metabolite concentrations reflect a balance between maximizing enzyme efficiency and minimizing total metabolite load. PLoS One. 2013;8.

[15] Noor E, Flamholz A, Liebermeister W, Bar-Even A, Milo R. A note on the kinetics of enzyme action: A decomposition that highlights thermodynamic effects. FEBS Letters. 2013;587:2772–2777.

[16] Schuster S, de Figueiredo LF, Schroeter A, Kaleta C. Combining metabolic pathway analysis with evolutionary game theory. Explaining the occurrence of low-yield pathways by an analytic optimization approach. Biosystems. 2011;105(2):147–153.

[17] Wessely F, Bart M, Guthke R, Li P, Schuster S, Kaleta C. Optimal regulatory strategies for metabolic pathways in Escherichia coli depending on protein costs. Molecular Systems Biology. 2011;7(515).

[18] Mori M, Marinari E, Martino AD. A yield-cost tradeoff governs Escherichia coli’s decision between fermentation and respiration in carbon-limited growth. biorxiv DOI:101101/113183. 2017;.

[19] Beg QK, Vazquez A, Ernst J, de Menezes MA, Bar-Joseph Z, Barabàsi AL, et al. Intracellular crowding defines the mode and sequence of substrate uptake by Escherichia coli and constrains its metabolic activity. Proceedings of the National Academy of Sciences. 2007;104(31):12663–12668.

[20] Müller S, Regensburger G, Steuer R. Resource allocation in metabolic networks: Kinetic optimization and approximations by FBA. Biochemical Society Transactions. 2015;43:1195–1200.

[21] Flamholz A, Noor E, Bar-Even A, Liebermeister W, Milo R. Glycolytic strategy as a tradeoff between energy yield and protein cost. Proceedings of the National Academy of Sciences. 2013;110(24):10039–10044.

[22] Wortel MT, Bosdriesz E, Teusink B, Bruggeman FJ. Evolutionary pressures on microbial metabolic strategies in the chemostat. Scientific Reports. 2016;6.

[23] Mori M, Hwa T, Martin OC, Martino AD, Marinari E. Constrained allocation flux balance analysis. PLoS Computational Biology. 2016;12(6):e1004913.

[24] Schuster S, Dandekar T, Fell DA. Detection of elementary flux modes in biochemical networks: A promising tool for pathway analysis and metabolic engineering. Trends in Biotechnology. 1999;17(2):53–60.

[25] Schuster S, Fell D, Dandekar T. A general definition of metabolic pathways useful for systematic organization and analysis of complex metabolic networks. Nature Biotechnology. 2000;18:326–332.

[26] Stelling J, Klamt S, Bettenbrock K, Schuster S, Gilles ED. Metabolic network structure determines key aspects of functionality and regulation. Nature. 2002;420:190–193.

[27] Jol SJ, Kümmel A, Terzer M, Stelling J, Heinemann M. System-level insights into yeast metabolism by thermodynamic analysis of elementary flux modes. PLoS Computational Biology. 2012;8(3):e1002415.

[28] Gerstl MP, Ruckerbauer DE, Mattanovich D, Jungreuthmayer C, Zanghellini J. Metabolomics integrated elementary flux mode analysis in large metabolic networks. Scientific Reports. 2015;5:Article number: 8930.

[29] Peres S, Jolicœur M, Moulin C, Dague P, Schuster S. How important is thermodynamics for identifying elementary flux modes? PLoS One. 2017;12(2):e0171440.

[30] Wortel MT, Peters H, Hulshof J, Teusink B, Bruggeman FJ. Metabolic states with maximal specific rate carry flux through an elementary flux mode. FEBS Journal. 2014;281(6):1547–1555.

[31] Müller S, Regensburger G, Steuer R. Enzyme allocation problems in kinetic metabolic networks: Optimal solutions are elementary flux modes. Journal of Theoretical Biology. 2014;347:182–190.

[32] Liebermeister W. Flux cost functions and the choice of metabolic fluxes. arXiv:180105742. 2018;.

[33] Holzhütter HG. The principle of flux minimization and its application to estimate stationary fluxes in metabolic networks. European Journal of Biochemistry. 2004;271(14):2905–2922.

[34] Noor E, Flamholz A, Bar-Even A, Davidi D, Milo R, Liebermeister W. The protein cost of metabolic fluxes: Prediction from enzymatic rate laws and cost minimization. PLoS Computational Biology. 2016;12(10):e1005167.

[35] Monk JM, Koza A, Campodonico MA, Machado D, Seoane JM, Palsson BO, et al. Multi-omics quantification of species variation of Escherichia coli links molecular features with strain phenotypes. Cell Systems. 2016;3(3):238–251.

[36] LaCroix AR, Sandberg ET, O’Brien JE, Utrilla J, Ebrahim A, Guzman IG, et al. Discovery of key mutations enabling rapid growth of Escherichia coli K-12 MG1655 on glucose minimal media using adaptive laboratory evolution. Applied and Environmental Microbiology. 2014;81:17–30.

[37] Long CP, Gonzalez JE, Feist AM, Palsson BO, Antoniewicz MR. Fast growth phenotype of E. coli K-12 from adaptive laboratory evolution does not require intracellular flux rewiring. Metabolic Engineering. 2017;44:100–107.

[38] Carlson R, Srienc F. Fundamental Escherichia coli biochemical pathways for biomass and energy production: Identification of reactions. Biotechnology and Bioengineering. 2004;85(1):1–19.

[39] Liebermeister W, Uhlendorf J, Klipp E. Modular rate laws for enzymatic reactions: Thermodynamics, elasticities, and implementation. Bioinformatics. 2010;26(12):1528–1534.

[40] Lubitz T, Schulz M, Klipp E, Liebermeister W. Parameter balancing for kinetic models of cell metabolism. Journal of Physical Chemistry B. 2010;114(49):16298–16303.

[41] van der Maaten PLJ, Hinton EG. Visualizing high-dimensional data using t-SNE. Journal of Machine Learning Research. 2008;9:2579–2605.

[42] van Rijsewijk BRBH, Nanchen A, Nallet S, Kleijn RJ, Sauer U. Large-scale 13C-flux analysis reveals distinct transcriptional control of respiratory and fermentative metabolism in Escherichia coli. Molecular Systems Biology. 2011;7(1):477.

[43] Maarleveld TR, Boele J, Bruggeman FJ, Teusink B. A data integration and visualization resource for the metabolic network of Synechocystis sp. PCC 6803. Plant Physiology. 2014; p. 113.

[44] Monod J. The growth of bacterial cultures. Annual Review of Microbiology. 1949;3(1):371–394.

[45] Shehata ET, Marr AG. Effect of nutrient concentration on the growth of Escherichia coli. Journal of Bacteriology. 1971;107(1):210.

[46] Alexeeva S, Hellingwerf KJ, de Mattos MJT. Quantitative assessment of oxygen availability: Perceived aerobiosis and its effect on flux distribution in the respiratory chain of Escherichia coli. Bacteriology. 2002;184(5):1402–1406.

[47] Bekker M, Kramer G, Hartog AF, Wagner MJ, de Koster CG, Hellingwerf KJ, et al. Changes in the redox state and composition of the quinone pool of Escherichia coli during aerobic batch-culture growth. Microbiology. 2007;153:1974–1980.

[48] López-Torrejón G, Jimènez-Vicente E, Buesa JM, Hernandez JA, Verma HK, Rubio LM. Expression of a functional oxygen-labile nitrogenase component in the mitochondrial matrix of aerobically grown yeast. Nature Communications. 2016;7.

[49] Basan M, Hui S, Okano H, Zhang Z, Shen Y, Williamson JR, et al. Overflow metabolism in Escherichia coli results from efficient proteome allocation. Nature. 2015;528:99.

[50] Fuhrer T, Fischer E, Sauer U. Experimental identification and quantification of glucose metabolism in seven bacterial species. Journal of Bacteriology. 2005;187:1581–1590.

[51] Andersen KB, von Meyenburg K. Are growth rates of Escherichia coli in batch cultures limited by respiration? Journal of Bacteriology. 1980;144(1):114–123.

[52] Szenk M, Dill KA, de Graff AMR. Why do fast-growing bacteria enter overflow metabolism? Testing the membrane real estate hypothesis. Cell Systems. 2017;5(2):95–104.

[53] Urbanczik R, Wagner C. An improved algorithm for stoichiometric network analysis: theory and applications. Bioinformatics. 2005;21(7):1203–1210.

[54] Klamt S, Regensburger G, Gerstl MP, Jungreuthmayer C, Schuster S, Mahadevan R, et al. From elementary flux modes to elementary flux vectors: Metabolic pathway analysis with arbitrary linear flux constraints. PLoS Computational Biology. 2017;13(4):e1005409.

[55] Beg KQ, Vazquez A, Ernst J, de Menezes AM, Bar-Joseph Z, Barabási AL, et al. Intracellular crowding defines the mode and sequence of substrate uptake by Escherichia coli and constrains its metabolic activity. Proceedings of the National Academy of Sciences. 2007;104:12663–12668.

[56] Goelzer A, Fromion V, Scorletti G. Cell design in bacteria as a convex optimization problem. Automatica. 2011;47(6):1210–1218.

[57] Goelzer A, Fromion V. Bacterial growth rate reflects a bottleneck in resource allocation. Biochimica et Biophysica Acta (BBA) - General Subjects. 2011;1810(10):978–988.

[58] O’Brien EJ, Lerman JA, Chang RL, Hyduke DR, Palsson BO. Genome-scale models of metabolism and gene expression extend and refine growth phenotype prediction. Molecular Systems Biology. 2013;9:693.

[59] Davidi D, Noor E, Liebermeister W, Bar-Even A, Flamholz A, Tummler K, et al. Global characterization of in vivo enzyme catalytic rates and their correspondence to in vitro kcat measurements. Proceedings of the National Academy of Sciences. 2016;113(12):3401–3406.

[60] Lerman JA, Hyduke DR, Latif H, Portnoy VA, Lewis NE, Orth JD, et al. In silico method for modelling metabolism and gene product expression at genome scale. Nature Communications. 2012;3:929.

[61] O’Brien EJ, Utrilla J, Palsson BO. Quantification and classification of E. coli proteome utilization and unused protein costs across environments. PLoS Computational Biology. 2016;12(6):e1004998.

[62] Dourado H, Maurino VG, Lercher MJ. Enzymes and substrates are balanced at minimal combined mass concentration in vivo. biorxiv DOI:101101/128009. 2017;.

[63] Valgepea K, Peebo K, Adamberg K, Vilu R. Lean-proteome strains–next step in metabolic engineering. Frontiers in Bioengineering and Biotechnology. 2015;3.

[64] Goel A, Eckhardt TH, Puri P, de Jong A, dos Santos FB, Giera M, et al. Protein costs do not explain evolution of metabolic strategies and regulation of ribosomal content: Does protein investment explain an anaerobic bacterial Crabtree effect? Molecular Microbiology. 2015;97(1):77–92.

[65] Teusink B, Passarge J, Reijenga CA, Esgalhado E, van der Weijden CC, Schepper M, et al. Can yeast glycolysis be understood in terms of in vitro kinetics of the constituent enzymes? Testing biochemistry. European Journal of Biochemistry. 2000;267:5313–5329.

[66] Scott M, Gunderson CW, Mateescu EM, Zhang Z, Hwa T. Interdependence of cell growth and gene expression: Origins and consequences. Science. 2010;330:1099.

[67] Valgepea K, Adamberg K, Seiman A, Vilu R. Escherichia coli achieves faster growth by increasing catalytic and translation rates of proteins. Molecular Biosystems. 2013;9(9):2344–23458.

